# SUMO-based expression and purification of dermcidin-derived DCD-1L, a human antimicrobial peptide, in *Escherichia coli*

**DOI:** 10.1101/343418

**Authors:** Ushanandini Mohanraj, Oona Kinnunen, Meryem Ecem Kaya, Aino Sesilja Aranko, Heli Viskari, Markus Linder

## Abstract

The dermcidin-derived peptide DCD-1L has a broad spectrum of antimicrobial activity over a wide pH range and in high salt concentrations. Thus, it offers a promising alternative to conventional antibiotics. Furthermore, it plays a role in wound healing, atopic dermatitis and acne vulgaris, indicating applications in cosmetic industries. Recently, dermcidin has been identified as a tumor marker improving cancer prognosis. Hence, large quantities of purified DCD-1L peptide are required to meet the needs of basic research and clinical trials. In the current study, we demonstrate SUMO-based heterologous DCD-1L production in *Escherichia coli*, followed by affinity chromatography purification. The SUMO tag is cleaved with SUMO-specific protease following purification, leaving free DCD-1L peptide without any additional amino acids. The mass of the peptide was further confirmed by MALDI-TOF-TOF analysis. Furthermore, the cleaved DCD-1L showed antimicrobial activity against the *E. coli* DH5 alpha test strain. The production and purification of DCD-1L using SUMO tag compare advantageously to other protocols previously described. Thus, the SUMO tag system enables large scale recombinant production of the antimicrobial peptide DCD-1L, which constitutes pharmaceutical and therapeutic potential as an alternative antibiotic.

## Introduction

New alternatives to antibiotics have received great attention due to an increasing threat of antibiotic resistance to commonly used antibiotics. Antimicrobial peptides emerge as a promising alternative due to their wide range of activity spectrum. Since the 70’s, several antimicrobial peptides from various species have been discovered. Antimicrobial peptides are known for their important role in natural self-defense mechanism for many species, including bacteria, fungi, and primates. Three important groups of human antimicrobial peptides that have been widely reported include defensins, histatins and cathelicidin. The third group comprises only one antimicrobial peptide, the cathelicidin LL-37. All three families of peptides are cationic in nature and they act by electrostatic interactions with the negatively charged phospholipid bilayer of pathogens (Smet et al., 2005). However, in 2001, a new anionic defense peptide called dermcidin was discovered in primates with no homology to other known AMPs (Schittek et al., 2001). Unlike human defensins and cathelicidins that are induced under inflammatory and injured conditions, dermcidin is constitutively expressed in human sweat (Rieg et al, 2004). The prevalent model for the structural basis of DCD-1L action is that it forms pores in the bacterial membrane leading to cell death (Paulmann et al, 2012).

Dermcidin is secreted to epidermal surface as a part of the first line of defense (Burian and Schittek, 2015). The dermcidin precursor is 110 amino acids long, and it includes a 19-aminoacid long signal peptide. Once the antimicrobial peptide precursor is secreted with sweat to the epidermal surface, the signal peptide is cleaved, and the precursor undergoes further proteolytic processing leading to formation of several dermcidin-derived peptides (Paulmann et al, 2012). Among these, DCD-1L is one of the most abundant dermcidin-derived peptides (Steffen et al., 2006). It is a 48-amino-acid long anionic peptide and it has an overall net charge of −2 with a cationic N-terminal region and an anionic C-terminal region (Paulmann et al., 2012, Steffen et al., 2006). DCD-1L is active against a wide range of pathogens including Gram-positive (*Staphylococcus aureus, Enterococcus faecalis, Staphylococcus epidermidis, Listeria monocytogenes*) and Gram-negative bacteria (*Escherichia coli, Pseudomonas putida, Salmonella typhimurium*) as well as *Candida albicans* (Steffen et al., 2006).

Dermcidin has evolved to survive and maintain its activity under a range of different pH and salt conditions that are characteristic of human sweat, which makes it possible to use it for a wide range of applications. It has been observed that wound healing is significantly impaired during reduced hBD-2 and DCD expression in burn wounds, which may explain the increased susceptibility to infection and sepsis in burn patients (Milner and Ortega, 1999). Furthermore, a significant reduction of DCD-derived peptides has been witnessed in the atopic dermatitis patients who encounter recurrent bacterial or viral infections and pronounced colonization with *Staphylococcus aureus* (Rieg et al., 2005). Also, the expression of DCD is downregulated in the sweat of patients with acne vulgaris, suggesting a deficiency in the constitutive innate defense (Nakano et al., 2015).

Due to the wide host range of dermcidin and its pharmaceutical and therapeutic potential, a cost-effective and scalable method for DCD-1L production is required. In the present work, we describe the successful cloning, expression and purification of DCD-1L using Smt3 as fusion partner without any additional residues and have demonstrated its activity. Employing SUMO tag is beneficial during primary purification step due to its positive effect on protein solubility. Further it also prevents the formation of inclusion bodies of the fusion proteins. SUMO is a highly conserved family of proteins in eukaryotes and is absent from prokaryotes (Malakhov et al., 2004). These proteins belong to a group of ubiquitin-like proteins due to their shared structural homology with ubiquitin. SUMO tags can be cleaved by using SUMO-specific proteases Ulp1 and Ulp2 in yeast, and SENP1 and SENP2 in humans (Yan et al., 2009). These proteases recognize SUMO through tertiary interactions and cleave it at the C-terminus (Malakhov et al., 2004). We have utilized Ulp1 enzyme in our work to cleave off the His6x-Smt3 tag that is used in expression and purification. The His6x tag in the N-terminus is used for purification with immobilized metal ion affinity chromatography columns designed for histidine-tagged proteins.

## Materials and Methods

### 2.1. Bacterial strains and plasmids

Chemically competent *Escherichia coli* TOP10 cells were used as host for cloning and *E. coli* T7 express cells were used for gene expression of His6x-Smt3-DCD-1L fusion constructs. *E. coli* was grown at 37 °C in LB media for cloning and expression, and kanamycin (50 µg/ml) was added to the media during growth of plasmid-containing strains. The vector pET28a(+) (carrying an N-terminal His6x tag and the gene for kanamycin resistance) was used for cloning and expression of the target genes. The vector allows for expression of exogenous proteins and peptides under the control of the T7lac promoter (Novagen, Madison, WI). Vector carrying the Smt3 tag and the pET28a(+) vector were kindly provided by Sesilja Aranko from Aalto University.

### 2.2. Cloning of the DCD-1L expression vector

Smt3 tag was amplified by PCR from with forward primer 5’- TAT<u>CATATG</u>GGATCGGACTCAGAAGTC-3’ (NdeI restriction site underlined) and reverse primer 5’-TGATCTCGAGTTA<u>GGATCCACC</u>AATCTGTTC-3’ (XhoI restriction site in bold; BamHI restriction site underlined). The PCR product was ligated into the vector pET28a(+) restricted with NdeI and XhoI restriction enzymes to obtain the plasmid pET28a(+)-Smt3.

The following DNA construct (for DCD-1L) was ordered from Integrated DNA Technologies (IDT) as a gBlock:

5’- AAC<u>GGATCC</u>AGCCTGCTGGAAAAAGGCCTGGATGGCGCGAAAAAAGCGGTGGG CGGCCTGGGCAAACTGGGCAAAGATGCGGTGGAAGATCTGGAAAGCGTGGGCAAAGGCGCGGTGCATGAT-3’ (BamHI restriction site underlined; XhoI restriction site in bold)

The DCD-1L gBlock from IDT was prepared according to the IDT instructions. DNA was dissolved in TE buffer (10 mM Tris; 0.1 mM EDTA; pH 8,0) at 10 ng/μl final concentration. Then the gBlock was ligated into the vector pET28a(+)-Smt3 restricted with BamHI (NEB) and XhoI (NEB) to obtain the plasmid pET28a(+)-Smt3-DCD- 1L (Figure 1)

**Figure 1.**
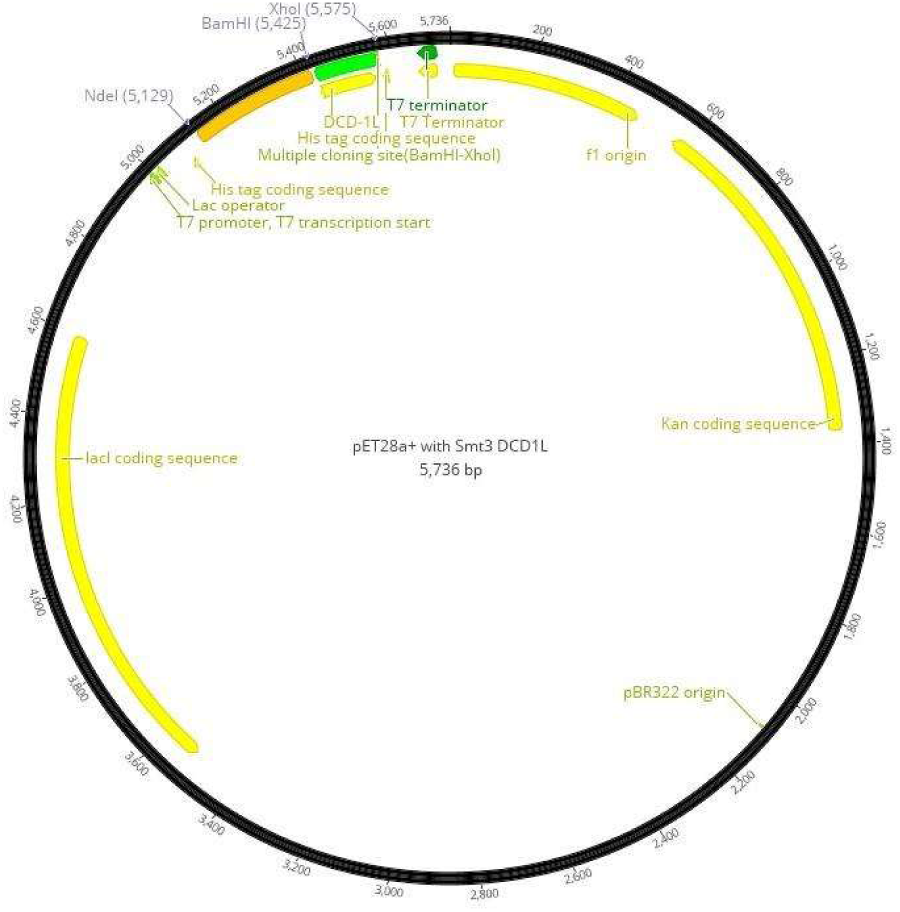
Plasmid map of Smt3-DCD-1L (Geneious version 10.1.3 [Kearse et al., 2012]).

Standard heat-shock transformation protocol was performed. Plates were incubated at 37 °C for over 12 hours before selecting kanamycin-resistant transformants for plasmid isolation, which was performed according to the manual of high copy number plasmid isolation in the Macherey-Nagel Nucleospin Plasmid kit. To verify that the isolated plasmids carried the desired insert, check-up digestion of the plasmids was performed using NdeI and XhoI restriction enzymes. Plasmids that carried an insert of the correct size were sent for sequencing at Eurofins Genomics, using the available sequencing primer T7 term (5’- CTAGTTATTGCTCAGCGGT-3’).

### 2.3. Small-scale expression and purification of DCD-1L

For small-scale production, T7 Express Competent *E. coli* were transformed with cloned plasmids by heat-shock transformation. 3-4 colonies were inoculated in 7 ml of LB-kanamycin (50 μg/ml working concentration). The cultures were incubated at 37 °C in shaking flasks. Expression of the construct was induced at an OD600 of ~0.5 with addition of IPTG at a final concentration of 0.5 mM. Cells were harvested 4 hours after the induction by centrifuging at 12,000 − g for 1 minute. The pellet was resuspended in 100 μl of ThermoFisher Scientific B-PER Bacterial Protein Extraction Reagent. After equilibrating the Qiagen Ni-NTA spin columns with NPI-10 buffer (50 mM NaPi, 300 mM NaCl, pH 8.0), protein purification was performed. Samples were loaded onto the spin columns and centrifuged at 1,600 rpm for 5 minutes, followed by washing the columns with NPI-20 buffer (50 mM NaPi, 300 mM NaCl, 30 mM imidazole, pH 8.0). Proteins were eluted in NPI-500 buffer (50 mM NaPi, 300 mM NaCl, 250 mM imidazole, pH 8.0). The used spin columns were soaked in 0.1 M EDTA solution and stored at 4 °C. Different flow-throughs and elution fractions were then analysed on 15% SDS-PAGE gel. The PageRuler^TM^ Prestained Protein Ladder (ThermoFisher) was used in this and other SDS-PAGE analyses. Gels were stained with Coomassie Blue and imaged with Bio-Rad Gel Doc^TM^ XR+ imager and Image Lab software (version 5.1). During purification, elution fractions were collected. Fractions containing the protein of interest were stored for further experiments.

### 2.4. Larger Scale expression and purification of DCD-1L

For larger scale expression, T7 Express Competent *E. coli* were transformed with cloned plasmids by heat-shock transformation. Kanamycin-resistant transformants were pre-cultured in LB medium with 50 μg/ml kanamycin at 30 °C overnight. The following day the pre-culture was diluted 1:100 with fresh LB-kanamycin medium and grown at 37 °C until OD600 value reached 0.6. Protein expression was induced with IPTG at a final concentration of 0.5 mM. Cells were harvested 4 hours after induction of protein expression by centrifuging in ThermoScientific Sorvall Lynx 4000 centrifuge at 5,000 − g for 10 minutes at 16 °C with the rotor F10-4−1000 LEX. Pellet was resuspended in Buffer A (50 mM NaPi, 300 mM NaCl, pH 8.0) and frozen in liquid nitrogen before storing at −20 °C.

Frozen cells were thawed in water bath before lysing them with EmulsiFlex-C3. Lysed cells were centrifuged in ThermoScientific Sorvall Lynx 4000 centrifuge at 18,000 − g for 30 minutes at 4 °C with the rotor F20-12−50 LEX. Supernatant was purified with GE Healthcare Life Sciences ÄKTA pure using the GE Healthcare Life Sciences HisTrap^TM^ FF crude 5 ml column. The proteins were eluted with Buffer B (50 mM NaPi, 300 mM NaCl, 250 mM imidazole, pH 8.0). Samples from different steps of protein expression and purification (non-induced, 2 hours after induction, 4 hours after induction, lysate, pellet, flow-through fractions from purification, eluates) were analysed on an SDS-PAGE to determine which fractions contained the protein of interest. Fractions containing protein of interest were pooled and stored at −20 °C.

To remove imidazole from purified proteins, buffer exchange and concentration steps were performed. Protein samples were concentrated with Sartorius Vivaspin 20 ultrafiltration centrifugal tubes (5,000 MWCO). The buffer was exchanged to sodium phosphate buffer (10mM, pH 7.4) such that the residual amount of Buffer B was ~5%. Protein samples were frozen in liquid nitrogen before storing at −20 °C. Concentration and final yield of DCD-1L were calculated from fusion protein concentration based on the hypothetical molecular mass ratio between the DCD-1L peptide and the fusion protein.

### 2.5. Ulp1 digestion for cleaving His6x and Smt3 tag from DCD-1L

Purified recombinant proteins containing His6x tag and Smt3 tag were cleaved with Ulp1 protease. 50 μl of recombinant protein mixture was incubated with 0.5 μl Ulp1 for 10-30 minutes at room temperature, after which the enzyme was heat inactivated for 10 minutes at 65 °C. Samples from cleaved and uncleaved proteins were analysed on 15% SDS-PAGE to verify the successful digestion.

### 2.6. Characterization of the purified recombinant peptide DCD-1L

Masses of purified peptides were identified with MALDI-TOF-TOF mass spectrometry. Salt was removed from Ulp1-digested DCD-1L peptide with ZipTip. Peptide samples and calibration standards were mixed with a final concentration of 10% acetic acid. Matrix was prepared for peptides by dissolving a pinch of α-cyano-4-hydroxycinnamic acid in a 7:3- mixture of deionized water and acetonitrile, and adding trifluoroacetic acid in a final concentration of 0.05%. Matrix was mixed thoroughly by vortexing for ~10 minutes. 1 μl of peptide sample or standard was mixed with 1 μl of the matrix on MALDI target plates. The measurement was performed with UltrafleXtream^TM^ Bruker MALDI-TOF-TOF mass spectrometer equipped with a 200-Hz smart-beam 1 lazer (337 nm, 4 ns pulse). Data collection was carried out by operating the instrument in positive ion mode controlled by the flex software package (FlexControl, FlexAnalysis). 5000 laser shots were accumulated per each spectrum in MS modes. Peptide calibration standard II (Bruker Daltonics) was used to calibrate the MS spectra.

### 2.7. Antimicrobial Activity of DCD-1L

Antimicrobial assays were performed to test the activity of DCD-1L against *E. coli* DH5 alpha strain. To obtain a standard curve for CFU/ml with respect to different OD600 values, the cells were grown to the desired OD600 values (0.1, 0.2, 0.3, 0.4, 0.5) and plated at 1:10^6^ dilution as described previously (Marc and Birgit, 2015). The plates were incubated at 37 °C for over 12 hours, and the CFUs corresponding to each OD600 value were calculated from the plates. The experimental *E. coli* cultures were grown at 37 °C until OD600 reached 0.05 (corresponding to 1.4−10^8^ CFU/ml). Cultures were incubated with antimicrobials (DCD-1L, LL-37, nisin and chloramphenicol at a concentration of 100 μg/ml) and control (sterile deionized water) at 37 °C with shaking for 40 minutes. After incubation, OD600 values of the cells were measured and corresponding CFU values were determined from the standard curve. The data are represented as the percentage of cells killed.

## 3. Results

### 3.1. Expression of DCD-1L peptides in bacteria

We aimed to produce and purify DCD-1L in *E. coli* using the SUMO fusion system. For His6x-Smt3-DCD-1L, the expression of the desired protein was observed through SDS-PAGE analysis after 2 hours and 4 hours of induction (Figure 2,3). In Figure 2, protein of interest can be observed in the eluate in lane 8. Furthermore, the protein was present in the lysate (Figure 2,3, lanes 4) and not in pellet (Figure 2,3, lanes 5), indicating good solubility of the fusion protein.

**Figure 2.**
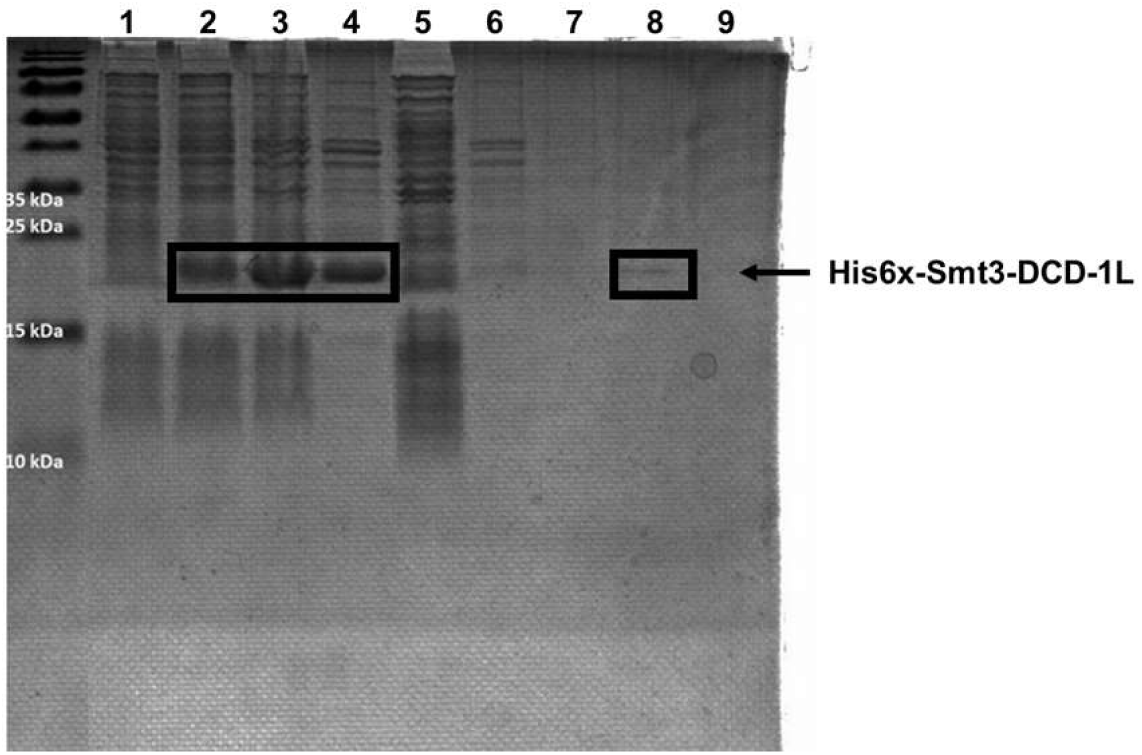
Analysis of His6x-Smt3-DCD-1L small-scale production. The molecular mass of His6x-Smt3-DCD-1L is 18.3 kDa and it is indicated in the figure with an arrow. Samples: 1. Smt3-DCD-1L non-induced, 2. Smt3-DCD-1L 2h after induction, 3. Smt3-DCD-1L 4h after induction, 4. Smt3-DCD-1L lysate, 5. Smt3-DCD-1L pellet, 6. Smt3-DCD-1L flow-through 1, 7. Smt3-DCD-1L flow-through 2, 8. Smt3-DCD-1L eluate 1, 9. Smt3-DCD-1L eluate 2.

**Figure 3.**
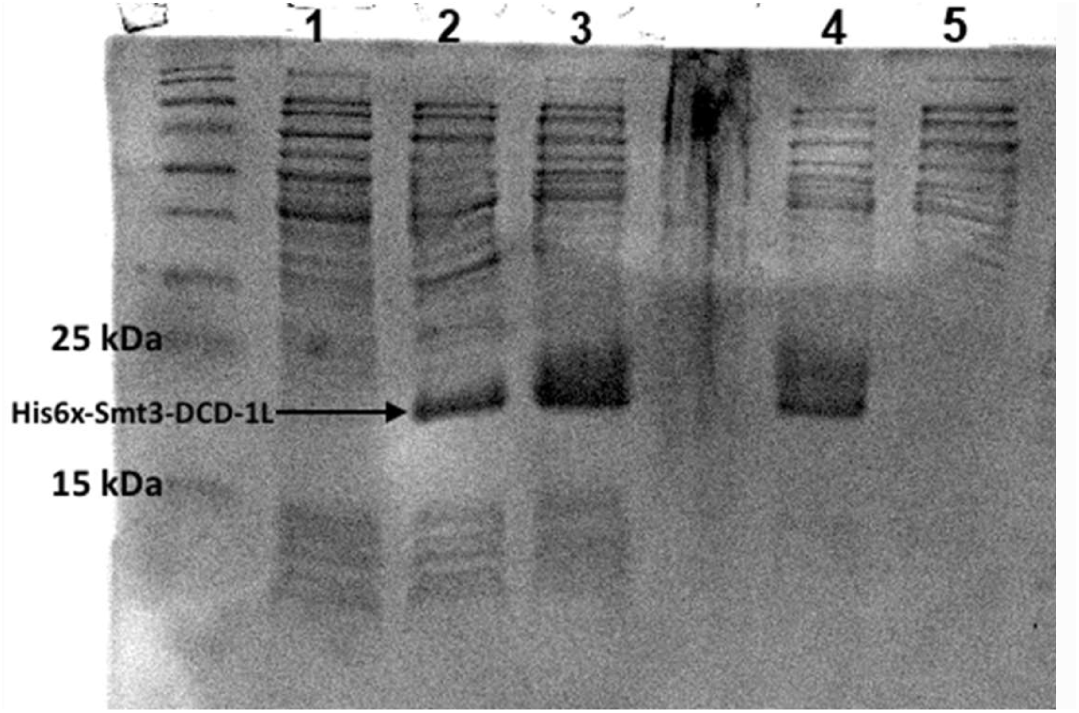
Analysis of His6x-Smt3-DCD-1L larger scale production. Samples: 1. Smt3-DCD-1L non-induced, 2. Smt3-DCD-1L 2h after induction, 3. Smt3-DCD-1L 4h after induction, 4. Smt3-DCD-1L lysate, 5. Smt3-DCD-1L pellet.

Detection of the intact fusion protein in the SDS-PAGE analysis provided evidence that the SUMO-peptide fusion was protected against cleavage by endogenous proteases. If cleavage had occurred during the purification step, the His6x-Smt3 tag (13.5 kDa) would be observed in the analysis of purified samples (Figure 2,3). Thus, it is clear that no endogenous cleavage interfered with the purification.

### 3.2. Purification of peptides

The His6x-Smt3 tag-based purification of the DCD-1L peptide was successful for small-scale and larger scale culture batches. In Figure 4, the protein of interest can be observed in the eluates, in lanes 1 to 6 corresponding to the molecular mass of His6x-Smt3-DCD-1L which is 18.3 kDa.

**Figure 4.**
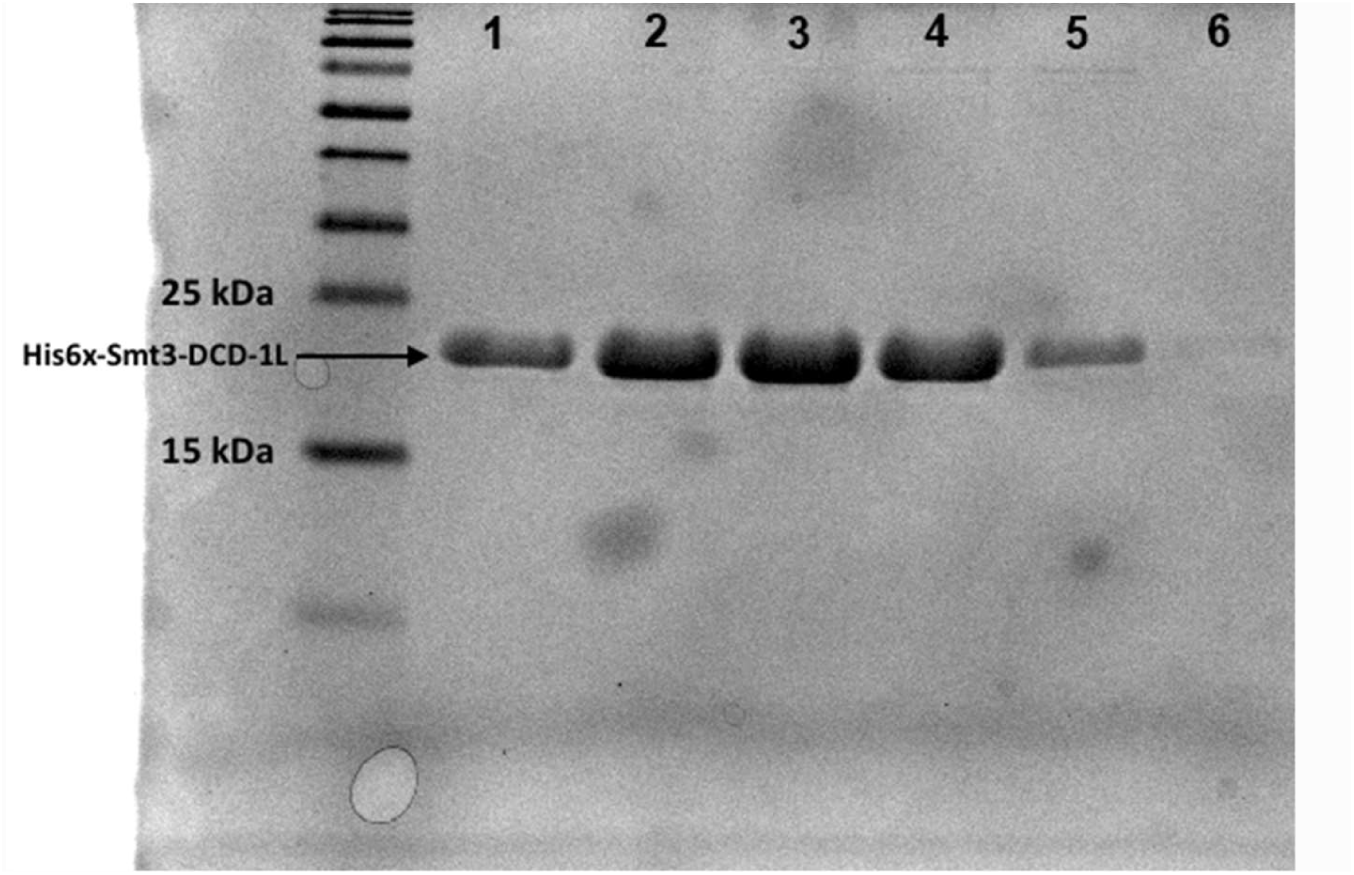
Analysis of the purified His6x-Smt3-DCD-1L. The molecular mass of His6x-Smt3-DCD-1L is 18.3 kDa. Samples: 1. Eluate fraction 1, 2. Eluate fraction 2, 3. Eluate fraction 3, 4. Eluate fraction 4, 5. Eluate fraction 5, 6. Eluate fraction 6.

From 500 ml culture, 25.47 mg of purified DCD-1L peptide was obtained after buffer exchange and concentration (Table 1). We anticipate that the construct and production system would likely require optimization to achieve higher yields.

**Table 1.** Table of final DCD-1L yield.

| Construct name | Culture volume [ml] | Purified protein volume [ml] | Concentration* [mg/ml] | Yield [mg] |
| --- | --- | --- | --- | --- |
| DCD-1L | 500 | 9 | 2.83 | 25.47 |

\*concentration of DCD-1L was calculated based on the fusion protein concentration according to the molecular masses of the peptide/fusion protein

### 3.3. Ulp1 enzyme digestion and SDS-PAGE

To determine a suitable reaction time for digestion of the fusion protein, the digestion reaction was incubated for different time intervals of 0 min, 5 min and 30 min. From Figure 5, it was observed that 5 min incubation already resulted in the digestion of proteins.

**Figure 5.**
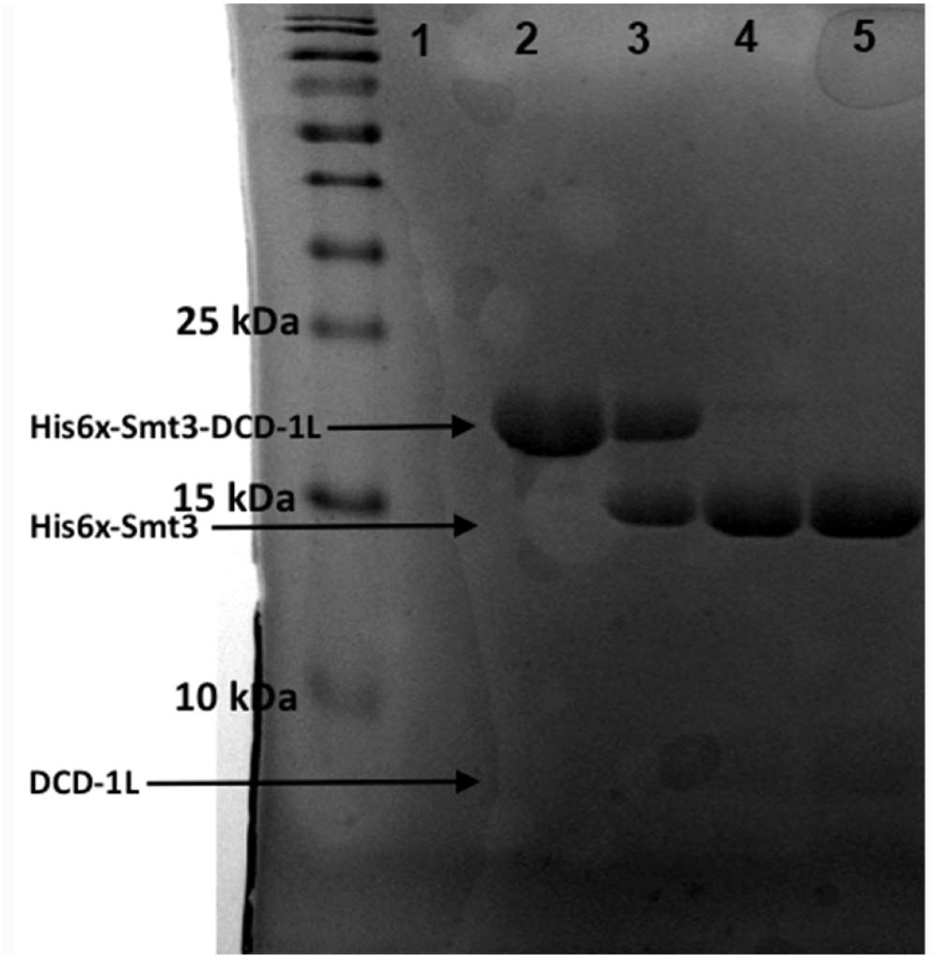
Digestion of His6x-Smt3-DCD-1L with Ulp1. Samples on the gel are: 1. Ulp1 (control), 2. His6x-Smt3-DCD-1L (undigested), 3. His6x-Smt3-DCD-1L + Ulp1 (digested for 1 minutes), 4. His6x-Smt3-DCD-1L + Ulp1 (digested for 5 minutes), 5. His6x-Smt3-DCD-1L + Ulp1 (digested for 30 minutes).

The molecular mass of the DCD-1L peptide is 4.82 kDa and the molecular mass of the undigested fusion protein His6x-Smt3-DCD-1L is 18.3kDa. The digested N-terminus containing the His6x-tag and the Smt3-tag is 13.5 kDa (Figure 5). Due to the small size of DCD-1L peptide, it is challenging to observe the corresponding bands on the SDS-PAGE gel. Instead, the band corresponding to the cleaved N-terminus containing His6x-Smt3 tag was clearly observed from the gel. The image further illustrates that the enzymatic digestion with Ulp1 is very rapid as the digested His6x-Smt3 tag can be clearly distinguished from the undigested protein within few minutes.

### 3.4. Identification of purified peptides

The identity of the peptides was confirmed by mass spectrometry. Following cleavage of expression tags from the fusion protein, DCD-1L peptide was analysed using mass spectrometry. A peak was observed at 4820.434 corresponding to the theoretical mass of DCD-1L peptide (Figure 6). A second peak was observed at 2410.626 and it corresponds to half the mass of the DCD-1L peptide. Given that the result from MALDI-TOF-TOF is mass/charge ratio and the overall charge of DCD-1L is −2, second peak is a further confirming result.

**Figure 6.**
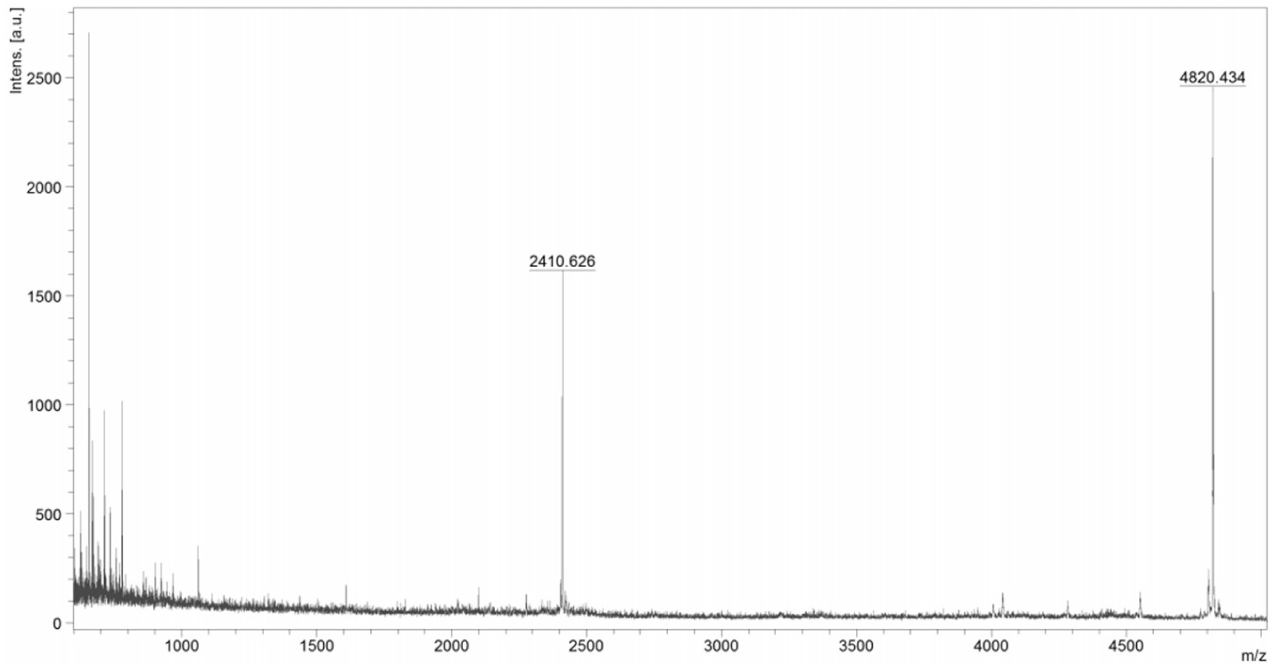
Image of mass spectrometry spectrum for DCD-1L with a peak around 4820.43 Da.

### 3.5. Antimicrobial assay

DCD-1L antibacterial activity along with other antimicrobials (chloramphenicol, nisin and LL-37) was tested against DH5 alpha strain of *E. coli*. Figure 7 illustrates that DCD-1L inhibited bacterial growth at a comparable level to other well-known antimicrobials, killing 70% of the cells, while nisin killed 64.28% and LL-37 57.85% of the cells.

**Figure 7.**
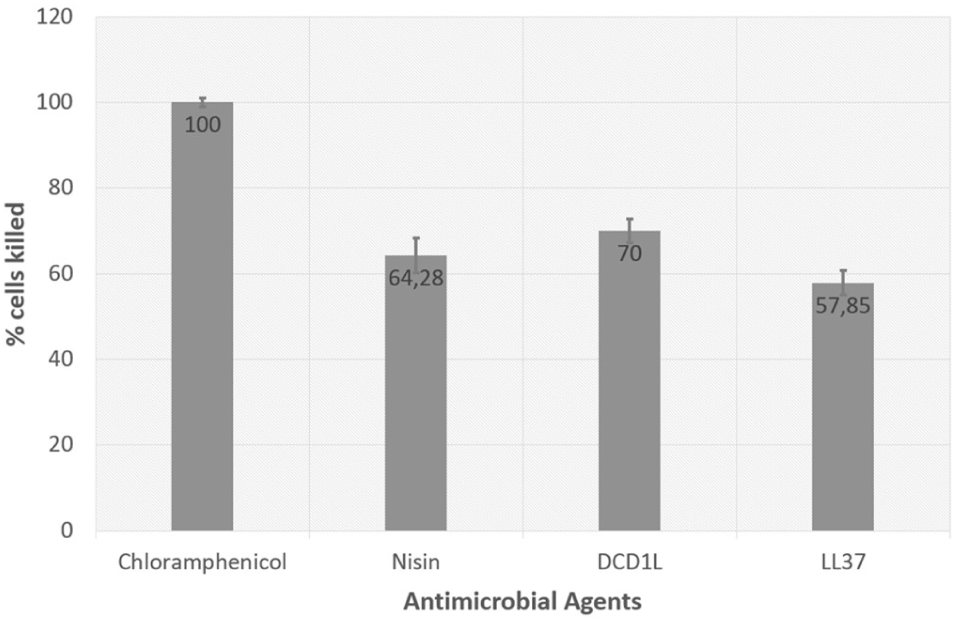
Antibacterial activity of DCD-1L along with Chloramphenicol, Nisin and LL-37 against E. coli DH5 alpha strain after 40 mins of incubation. Antimicrobial activity is represented in terms of percentage of cells killed.

## Discussion

Despite being discovered in 2001, only few production systems for dermcidin have been reported in the literature. The few methods that have been reported for the production and purification of dermcidin are based on the chemical and enzymatic routes to cleave the peptide from its fusion partner. Previously a CNBr cleavage-based method has been reported for the recombinant production of a 48-amino-acid dermcidin variant with a C-terminal homoserine lactone (DCD-1Hsl) in *Escherichia coli*, expressed in the form of inclusion bodies (Cipáková et al., 2006). Cyanogen bromide, which cleaves C-terminally after methionine, is used extensively, yet it is inefficient in its cleavage (Haught et al., 1998). The presence of methionine within the sequence of many antimicrobial peptides limits its application. In addition, methionines in the carrier or linker region make subsequent purification complicated and less efficient. In the case of the expression of the SUMO-peptide fusion, the fusion protein does not have to be forced into insoluble fractions. On an industrial scale, this could be costly due to the requirement for urea and guanidium chloride to solubilize the inclusion bodies (Lee et al., 2000). Moreover, as observed from our current study, with SUMO system the fusion protein remained in the soluble fraction.

In another study, recombinant MDCD-1L was expressed as an intein fusion protein in *E. coli*, and then purified by affinity chromatography using chitin beads (Hong et al, 2010). Intein system eliminates the need for exogenous proteases or chemicals that are usually required for tag removal and allows the target peptide to be purified through relatively simple procedures. However, the large size of the intein/chitin domain is a drawback for peptide yield. In addition, uncontrolled autocleavage of the intein fusions even at low levels could be a serious problem as the released antimicrobial peptides can cause fatal effects through direct interaction with their intracellular targets (Díaz et al., 2012). SUMO has been shown to improve folding and its small size (11.2 kDa) allows a relatively high peptide-to-carrier ratio, which favours peptide yield (Li, 2009). In our study, we obtained a yield of 25.47 mg per 500 ml batch of *E. coli* culture. The urea-resistant property of SUMO is a big advantage as the presence of a small amount of urea can prevent protein aggregation and loosen compact structures, which favours the cleavage (Li, 2011).

A highly specific SUMO protease facilitates efficient release of the peptide of interest and leaves no unwanted amino acids at the N-terminus of the peptide after cleavage (Li, 2011). From our mass spectrometry results after Ulp1 digestion, we observed no scar on the peptide. Another important advantage is that this protease is produced cheaply using the T7 driven pET system and can be easily purified with Ni-NTA affinity chromatography similar to the DCD-1L production system and hence, enzymes can be produced in large quantities with relative easiness. In our study, a low amount of enzyme was used for complete cleavage reducing the cost of peptide production. Another study reported the use of factor Xa protease to cleave the fusion protein to produce recombinant DCD-1L (Lai et al., 2005). Comparing with SUMO, factor Xa is more expensive and sensitive to pH and chaotropes.

In our study, we observed antimicrobial activity with concentration of 100 μg/ml. This is contradictory to the previous studies, where the minimum inhibitory concentration of DCD-1L was reported to be 10 μg/ml (Schittek et al., 2001, Steffen et al., 2006). One of the possible reasons for such difference is the *E. coli* test strain used in this study, which was a cloning strain unlike the ones used in literature. Further comparative antimicrobial activity assays with different bacterial strains need to be performed. One of the current limitations of the study is that data on the activity against other bacterial species are lacking. Thus, the activity of DCD-1L should be studied further against pathogens including *Streptococcus* and *Propionibacterium* acnes. Another limitation of the study is that the cleaved tags and the heat inactivated Ulp1 enzyme residues were not removed from our final purified protein products using reversed-phase HPLC before investigating the antimicrobial activity. This limits our ability to obtain reliable results on the antimicrobial activity of our produced peptide.

Based on different studies it seems to be contradictory whether the presence of salt affects the antimicrobial activity of DCD-1L. The presence of divalent cations (especially *Zn*^2+^, but also *Mg*^2+^, *Ca*^2+^) and monovalent cations (*Na*^+^ has reportedly increased the activity of DCD-1L. It seems to be important for the structure formation of the DCD-1L complex in the bacterial membrane (Paulmann et al., 2012). However, in another study the addition of high concentrations (100 mM) of NaCl to the buffer was shown to decrease the activity against *E. coli* (Schittek et al., 2001). Thus, we think that the potential contribution of mono- and divalent cations to the antimicrobial activity of DCD-1L could be studied further. Hence, further studies need to be carried out to analyse the activity of DCD-1L in different salt and pH conditions.

## Conclusions

In summary, this work presents an expression and purification method that can be applied to the production of the human anionic antimicrobial peptide DCD-1L in bacteria on large scale. Since DCD-1L is reported to have a wide range of antimicrobial activity and implications in cosmetic industry, we can conclude that this production and purification approach provides a powerful tool for mass production of biologically active DCD-1L in industries. Also, in the future, engineering DCD-1L variants with improved antimicrobial properties have a great potential as antibiotics.

## Acknowledgements

We are grateful to Pezhman Mohammadi at Aalto University for helping with MALDI-TOF-TOF mass spectrometry analysis of the peptides and Prof. Per Erik Saris for providing us with valuable inputs about antimicrobial assays. We would like to thank Maisa Vuorte, Meo Ekroos, Matilda Tuure and Michal Pasik from Aalto University and Eveliina Karjalainen and Jenny Mujunen from University of Helsinki for their kind assistance and support in the lab experiments.

## Ethics Statement

N/A

## Data Availability

All relevant data are within the paper.

